# Genome-wide gene birth-death dynamics are associated with diet breadth variation in Lepidoptera

**DOI:** 10.1101/2023.04.11.536384

**Authors:** Hanna Dort, Wouter van der Bijl, Niklas Wahlberg, Sören Nylin, Christopher W. Wheat

## Abstract

Comparative analyses of gene birth-death dynamics (GBDD) have the potential to reveal gene families that played an important role in the evolution of morphological, behavioral, or physiological variation. Here, we used whole genomes of 30 species of butterflies and moths to identify GBDD among the Lepidoptera that are associated with specialist or generalist feeding strategies. Our work advances this field by using a uniform set of annotated proteins for all genomes, investigating associations while correcting for phylogeny, and assessing all gene familes rather than *a priori* subsets. We discovered that the sizes of several important gene families (e.g., those associated with pesticide resistance, xenobiotic detoxification, and/or protein digestion) are significantly correlated with diet breadth. We also found 12 gene families showing significant shifts in GBDD at the butterfly (Papilionoidea) crown node, the most notable of which was a family of pheromone receptors that underwent a contraction potentially linked with a shift to visual-based mate recognition. Our findings highlight the importance of uniform annotations, phylogenetic corrections and unbiased gene family analyses in generating a list of candidate genes that warrant further exploration.

**Significance Statement:** Gene duplications and gene deaths are important for the development of novel traits, but their study is often complicated by methodological issues, such as input data coming from different sources with different biases. Here, we address many of these issues in an analysis of gene duplication and death across 30 species of moths by using standardized, genome-wide input data. Importantly, our analysis uses a model that can correlate gene family sizes with any quantitative trait, while also accounting for relatedness between species. Using this model, we provide novel evolutionary insights by showing that the sizes of several important gene families are correlated with increases or decreases in the number of host plant orders that Lepidoptera eat.

## Introduction

Understanding the origins of novel traits is central to the study of evolution. At a genomic level, the influential work of Ohno (1970) convincingly argued that one route for the generation of novel traits was through novel genes, most often arising via gene duplication events. Following a duplication event, new gene copies can quickly accumulate mutations due to relaxed selection pressure. Although mutation usually results in gene copies becoming non-functional, such variation also allows for novel traits to develop, via gene neofunctionalization or specialization (for a review of these terms, see Lynch & Conery 2000; Lynch 2007). Novelty can also evolve through gene death or loss of function, as this pathway is often adaptive in the context of response to environmental change (Albalat & Cañestro 2016; Monroe et al. 2021). Thus, as evolutionary biologists work to find genes that cause novel traits, the study of gene birth and death dynamics (GBDD) is a powerful approach to be added to the comparative study of the regulatory and coding regions of genes (Sackton et al. 2019).

Due to their extensive natural history record and their early use in formulating coevolutionary theory (Ehrlich & Raven 1964), the Lepidoptera are often used as a model to explore links between GBDD and specific phenotypes of interest. For example, GBDD have been assessed for Lepidoptera in the contexts of hostplant detoxification (Fischer et al. 2008; Feyereisen 2011; Edger et al. 2015; Calla et al. 2017), hostplant specialization (Suzuki et al. 2018), hostplant identification (Briscoe et al. 2013), and the development of color vision (Sondhi et al. 2021). In most of these studies, researchers investigated differences in the copy number of genes in pre-determined gene families and ultimately found that birth events at both the exon and gene level were associated with key innovations. Gene death events have often been ignored in the insect literature, although they too can be associated with major adaptations, such as the evolution of BT pesticide resistance in *Helicoverpa* (Yang et al. 2006), a transition to visual mate recognition in Papilionoidea (Vogt et al. 2015), and a shift to herbivory within Drosophilidae through the loss of odorant receptors (Goldman-Huertas et al. 2015). In any case, the tendency to consider GBDD only in gene families with hypothesized importance has resulted in genome-wide GBDD remaining largely unexplored, though we note an excellent analysis of GBDD patterns across Arthropoda than included Lepidoptera species (Thomas et al. 2020). Thus, many important gene duplication and death events that are associated with diverse adaptations have yet to be discovered via unbiased, genome-wide comparative analyses of GBDD.

Insightful comparative analysis of GBDD relies upon several important factors, from high quality protein sets, to a reliable phylogenetic context for analysis. While such studies are providing insight into GBDD among diverse clades, there are a growing number of concerns. First, the vast majority of GBDD analyses use publicly available published genome annotations as analysis inputs. Since each genome is likely annotated using slightly different approaches, especially in the annotation of protein isoforms, the resulting annotation files are very heterogeneous and can dramatically skew estimates of GBDD, especially when used without any isoform filtering. Recent analysis of the effects of annotation heterogeneity on estimates of lineage-specific genes found the problem both widespread across the tree of life, and significant, inflating estimates upwards of 15-fold (Weisman et al. 2022). Simply put, these important findings call into question well over a decade’s worth of GBDD literature. Second, GBDD analyses typically use a time calibrated phylogenetic tree to identify specific nodes where births or deaths occurred, as well as estimate their rates. Unfortunately, most studies fail to disclose whether their analysis assumed that all gene families were present at the root of the tree (the default setting in CAFE), which will systematically over-estimate gene loss when some families are simply unique to a subset of taxa. Further, the vast majority of such studies estimate both the phylogenetic relationships, and the timing of divergence using genomic data from the species being studied. However, such datasets are generally very poor in taxon sampling, which can lead to biases in estimated phylogenetic relationships as well as times of divergence (Wheat and Wahlberg 2013). Detailed phylogenetic analyses along with robust estimates of times of divergence are available for most groups that have reference genomes, and this information can be used in GBDD analyses. Finally, investigations of GBDD being associated with phenotypes must take into account the non-independent relationships among the studied taxa (i.e. their shared evolutionary history). A failure to do so can easily generate significant, yet meaningless, correlations that will cloud the literature.

Here, we perform genome-wide GBDD analyses on 30 diverse species of Lepidoptera and investigate if the sizes of any gene families can be correlated with diet breadth along a specialist-generalist axis. This question is of evolutionary and ecological interest because generalist feeding is thought to reflect the mechanisms by which specialist lineages can colonize new plant species (Janz & Nylin 2008; Nylin et al. 2014), and shifting between these strategies is relatively common for Lepidoptera over evolutionary time (Nosil 2002; Braga et al. 2018). Although several previous studies have tried to quantify links between diet breadth and gene family expansions/contractions (e.g. Calla et al. 2017; Suzuki et al. 2018; Breeschoten et al. 2022), their scope was limited by only looking at gene families linked with detoxification and host preference, rather than investigating all genes in the genome. Unfortunately, several aspects of these studies’ methods also complicate interpretation of their findings, such as only comparing qualitative groups (i.e. generalists vs. specialists), using different genome annotation methods for different species, not filtering out isoforms from datasets, and/or failing to take into account the effects of shared ancestry among species in their analyses.

To address these issues, we used a standardized approach to generate protein annotations and filter protein sets for all species in our analyses, and we based our phylogenetic tree upon existing literature for the most accurate relationships and ages. We then quantified overall rates of gene duplication and death and investigated whether there were any significant gene family expansions or contractions associated with the origin of butterflies (i.e., at the Papilionoidea crown node). Finally, we explored whether the size of any gene families were correlated with species’ diet breadth, while taking into account the phylogenetic relationships among species, allowing for both positive and negative correlations. A functional assessment of gene families exhibiting significant correlations with diet breadth revealed associations with digestion, xenobiotic detoxification, and egg development, suggesting that there may indeed be key genetic differences between specialist and generalist Lepidoptera that warrant further study.

## Results & Discussion

### Genomes and *de novo* annotations were of high quality

Our dataset of 30 published Lepidopteran genomes covered eight superfamilies within Ditrysia (Tortricoidea, Pyraloidea, Noctuoidea, Papilionoidea, Bombycoidea, Gelechioidea, Geometroidea, and Yponomeutoidea), and 15 families in total (see Figure 1; Table S1). The number of host plant orders per species ranged from 1 to at least 6 (Figure 1). Genome quality overall, as assessed by completeness, was good, with each genome having on average 93.8% of Lepidoptera BUSCO genes complete and single copy in their genomes (Table S1). In order to obtain a uniform set of annotations for each species, we used the BRAKER2 pipeline to generate a new, standardized annotation for each species. An average of 24,836 genes were predicted per species, which was reduced to 19,948 after filtering to the single longest isoform per locus (Table S2). These had a good overall completeness, containing an average of 4697 of 5286 (88.9%) of Lepidoptera BUSCO genes in complete, single copies following filtering to remove isoforms and non-coding genes (Table S3).

**Figure 1.**
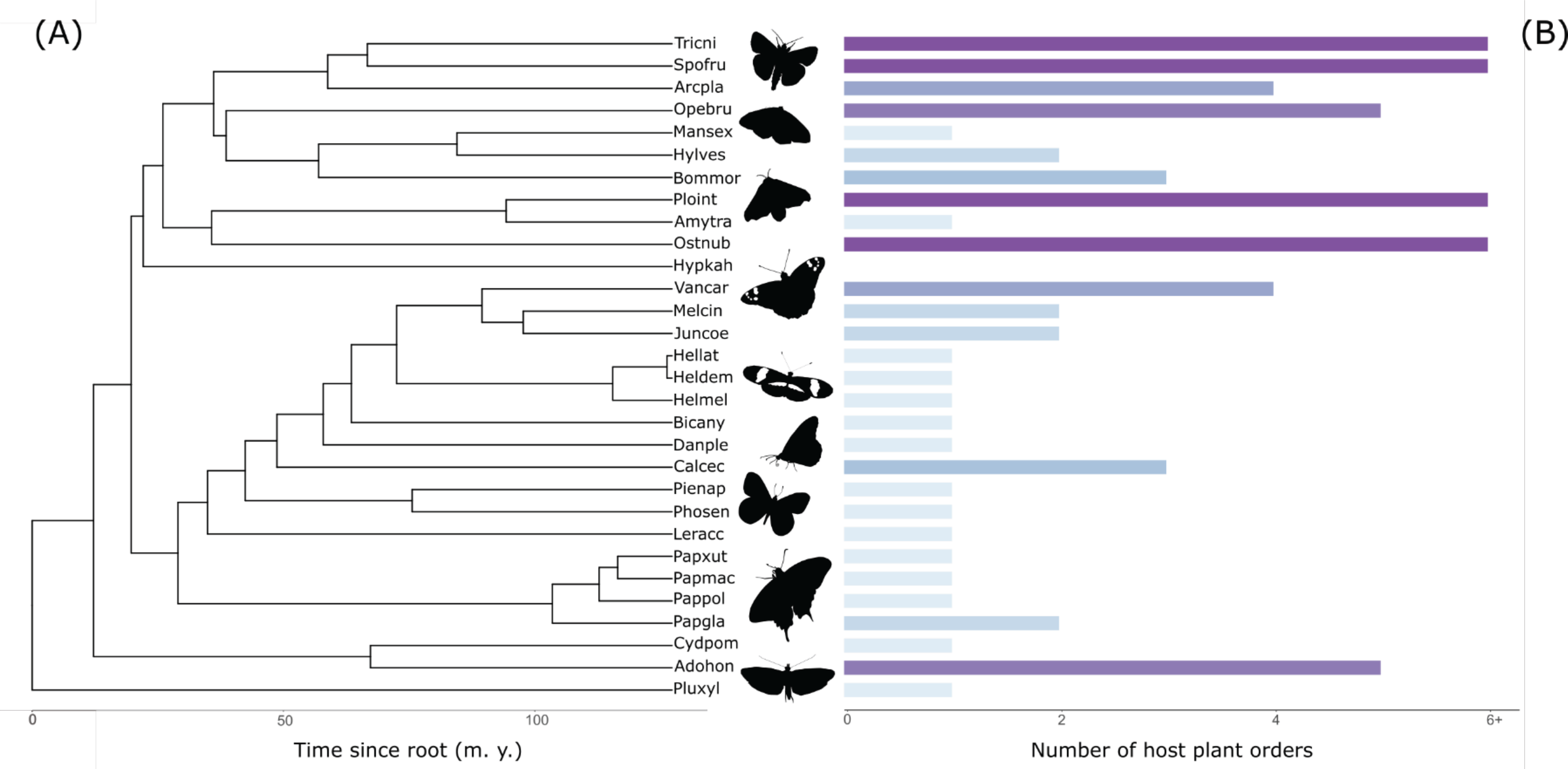
(A) A time-calibrated phylogeny of all species (N=30) used in our CAFE analyses. (B) Per-species counts of the number of host plant orders identified as being commonly used by larvae. For lists of these host plant orders, see Table S8.

For comparison, the average percentage of complete, single copy BUSCO genes found in the unfiltered, native protein set was 80.92% (Table S4), which is nearly 10% lower than our BRAKER annotations. The average percentage of duplicated BUSCO genes per species was higher in the unfiltered, native protein set (10.3%, Table S4) than in the filtered de novo protein set (3.2%, Table S3), likely due to the presence of isoforms in the native set.

### Filtering steps reduced gene counts for both native and *de novo* annotations

Because many previous GBDD studies have relied on datasets made from annotations or protein sets downloaded alongside reference genomes, we were interested in quantifying the effects basic filtering steps had on these “native” datasets and comparing them to the effects of filtering steps on our *de novo* dataset. Regardless of annotation source, final protein counts were consistently reduced by filtering datasets to only include single isoforms of genes with correct start/stop codons (Figure 2A; 2B). Although there was no clear trend of one annotation source being more effected than the other by filtering (Figure 2D), our filtered *de novo* annotations from BRAKER2 resulted in significantly higher per-species protein counts than filtered native annotations (Paired t-test: t = 12.469, df = 18, p-value = 2.714e-10; Figure 2C).

**Figure 2.**
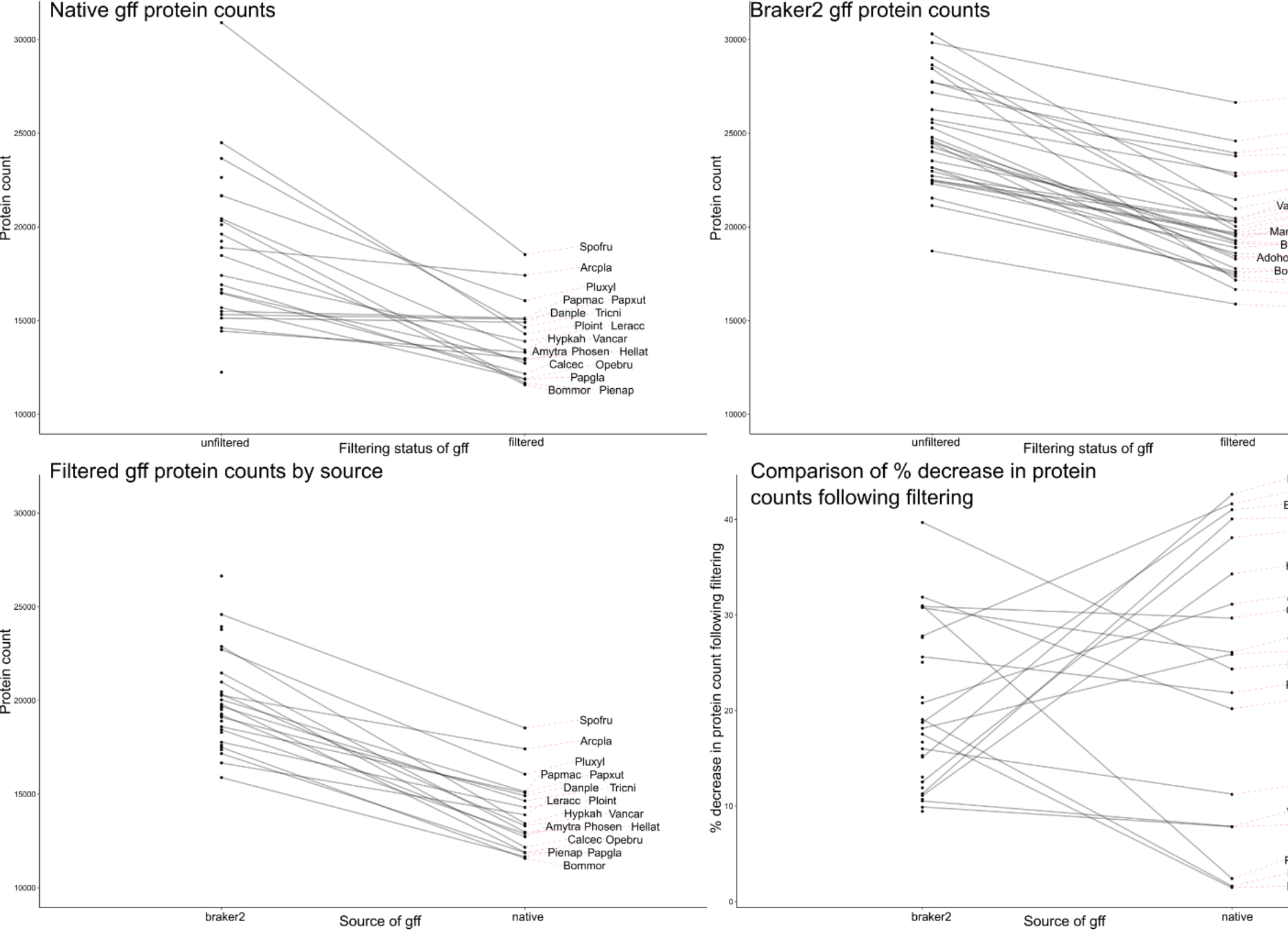
A comparison of the effects of filtering on predicted protein counts for both native and *de novo* (BRAKER2) annotations. For full species names, see Table S1. (A) Decrease in total protein count per species following filtering of native annotation files. Note that only 19 of 30 species included in this study had locatable native annotation files (e.g. a GFF file); as such, not all species have filtered native annotations. (B) Decrease in total protein count per species following filtering of *de novo* annotation files. (C) Comparison of total protein counts between filtered *de novo* and filtered native annotations for each species. (D) Per-species comparison of the effect (% decrease in protein counts) of filtering for *de novo* versus native annotations.

### Compared to *de novo* protein sets, native protein sets had inflated gene family sizes

For the 19 species with locatable native annotation files, filtering native annotations so that only proteins with correct start and stop codons were retained decreased protein counts by an average of 12.8% (Table S5). Additionally, filtering annotations to remove all but the longest isoforms for each gene decreased predicted protein counts by values ranging from 0 to 12,418 genes (Table S5). Together, these filtering results suggested that using native protein sets without additional modification would be inappropriate for GBDD analyses and led us to predict that unfiltered native protein sets would have greatly inflated gene family sizes, compared to our BRAKER2 protein sets, which were filtered with a standardized pipeline.

To test the above prediction, we ran all BRAKER2 protein sets and all raw, native protein sets through OrthoFinder in two separate runs. The output of these runs included estimates for, e.g., total gene counts and median number of genes per assigned orthologous gene cluster, or orthogroup (Table S5). Using the native annotation set, there were fewer orthogroups compared to the BRAKER2 set (Table 1; Table S6), though each native orthogroup had more than twice as many members (median gene count: 21.0) as the BRAKER2 set (median gene count: 9.0).

**Table 1.** A selection of results from two Orthofinder runs. The first run reported (filtered de novo protein set) was performed using full protein sets from 30 Lepidoptera species, which were created with a standardized genome annotation and filtering pipeline. The second run was performed using the protein sets provided with reference genome assemblies (native sets) with no further filtering occurring.

|  | Filtered, *de novo* protein set | Raw native protein set |
| --- | --- | --- |
| Total number of genes | 598 466 | 572 004 |
| Total number of orthogroups | 27 396 | 20 993 |
| Orthogroups present in all 30 species | 4118 | 3351 |
| Percent of genes assigned to an orthogroup | 90.9 | 93.7 |
| Mean number of genes per orthogroup | 19.9 | 25.5 |
| Median number of genes per orthogroup | 9.0 | 21.0 |
| Percent of genes in a species-specific orthogroup | 2.1 | 3.1 |
| Percent of orthogroups present in all 30 species | 15.0 | 16.0 |

Thus, the decisions made when cleaning and filtering protein sets (or failing to do so) could have unintended downstream effects for comparative genomics analyses. A failure to filter out isoforms will inflate gene birth rates in GBDD analyses as CAFE will interpret two isoforms of the same gene as genes born from a duplication event. Failing to use a standardized genome annotation approach for all species can further inflate the number of lineage-specific genes, as previously reported by Weisman et al. (2022). Highlighting these issues, in our comparison of native versus *de novo* generated protein sets, we found that native annotations had a much higher number of genes per orthogroup, and a marginally higher percentage of species-specific orthogroups (Table 1, Table S6). Because these differences could be due to a bioinformatic (i.e. poor filtering) rather than a biological origin, results based upon similarly non-filtered data should be viewed with caution.

### Gene birth rate surpasses gene death rate, provided gene families exist at the tree root

We first analyzed gene birth and death rates over the entire phylogeny with the assumption that gene birth rate was equal to gene death rate and for gene families that were present in the root ancestor (Table 2). By removing the assumption that gene duplication rate was equal to gene death rate and repeating this analysis, we found that model scores were consistently improved, indicating the importance of relaxing this simplifying assumption (Table S7). Ultimately, we found the average tree-wide rate of gene duplication (λ = 0.0034 duplications/gene/million years) was greater than the corresponding rate of gene death (μ = 0.0012 deaths/gene/million years) (Table 2).

**Table 2.** Estimations of tree-wide gene duplication and gene death rates (per gene per million years) calculated by CAFE v.4. Values reported here are averages from sets of three runs per model setting. For a full table of results, including model scores, see Table S7

| Model setting | lambda<br>(duplication rate) | mu<br>(death rate) |
| --- | --- | --- |
| lambda = mu | 0.002416423 | NA |
| lambda ≠ mu | 0.003410306 | 0.001164384 |

These results are perhaps surprising, as a previous study of tree-wide gene duplication and death rates for Lepidoptera estimated that gene death rate was roughly double gene duplication rate (μ = 0.0032 vs λ = 0.0015; Breeschoten et al. 2022). In addition to differences in annotation heterogeneity and filtering steps (Breeschoten et al. used raw, native protein sets without isoform filtering), our seemingly opposite results are likely also influenced by our choosing to only calculate rates for gene families present at the root of the phylogeny. It is our understanding that failing to perform this filtering step (i.e., to not use the –filter option in CAFE) would result in a greatly inflated gene death rate, as CAFE v.4 by default assumes all gene families in an input existed at the tree root. Thus, a gene family that originated recently in one species would be modelled as having died out in all other species. In the latest version of CAFE, this filter is now default (i.e. gene families not found at the root are excluded from estimates). We would, however, like to note that only including families present at the tree root also creates a bias because this set of genes might be under selection for their retention.

### GBDD analysis at the Papilionoidea crown node reveals a significant loss of odorant receptors, but no excess in births or deaths compared to other nodes

Given the range of traits that differentiate butterflies from moths, we expected there to be strong GBDD for traits associated with the crown node of Papilionoidea that distinguish butterflies from other Lepidoptera (e.g., genes responsible for diurnal behavior (Kawahara et al. 2018), color vision (Briscoe et al. 2013), or wing color patterns (Nijhout 2001). We explored butterfly GBDD associated with the crown node of Papilionoidea, which is approximately 100 million years old (Chazot et al. 2019), by investigating whether there were any major gene family expansions or contractions occurring on the branch leading this node in our BRAKER2 protein dataset.

We found no evidence of an excess of significant gene family expansions or contractions at the Papilionoidea crown node, as compared to other nodes (Fig. S1). There are many potential explanations for this observation. First, any necessary adaptations could have been via regulatory change rather than via copy number change. Second, changes in the life history strategies of early butterflies may not have been due to novel adaptations, but rather due to ecological opportunity (see Araujo et al. 2015). Third, exaptation of existing gene families might have been sufficient, obviating the need for dramatic changes in gene counts at the Papilionoidea crown (e.g. Gould & Vrba 1982). The most likely explanation, however, is that the key innovations we associate with butterflies stem from gene family expansions and contractions at a subfamily or genus level, as many major life history changes (such as dietary specialization) often occur at those smaller scales. Such events may leave complicated signatures at deeper evolutionary time scales.

However, we identified an exciting, significant contraction in a family of 7tm odorant receptors in our BRAKER2 dataset (Table 3; Figure 3).

**Figure 3.**
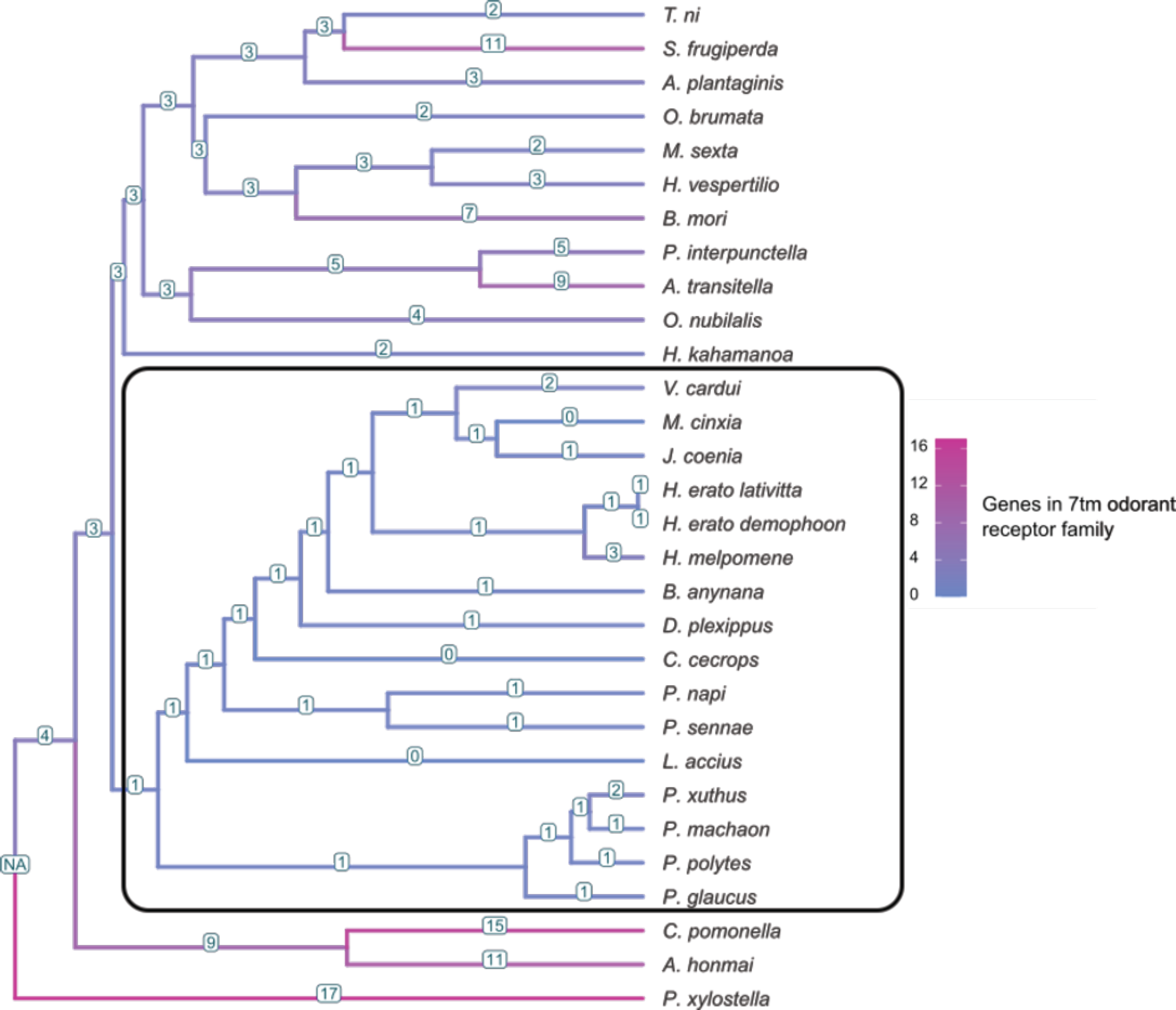
Estimated counts of genes in the 7tm odorant receptor (putative pheromone receptor) family identified as rapidly contracting at the crown of Papilionoidea. The Papilionoidea clade is within the black box. Note that all non-butterfly moths have several odorant receptor genes in this family, but most butterflies have one copy. Counts were taken from a root-filtered, *de novo* generated input where gene birth rate was not assumed to equal gene death rate.

**Table 3.** Functional annotations and detected size changes (in # genes) of gene families found to be significantly expanding or contracting at the root node of the Papillionoidea.

| ID | Change in # genes | eggNOG functional annotation(s) |
| --- | --- | --- |
| 23 | 4 | zinc finger; phosphatidyl serine synthase |
| 478 | 2 | DNA polymerase |
| 40 | -5 | TRAFAC class kda protein, type-1-like; mysosin-kinesin ATPase superfamily |
| 69 | -3 | zinc finger |
| 80 | -5 | reverse transcriptase |
| 89 | -5 | reverse transcriptase |
| 131 | -2 | cuticle protein |
| 249 | -3 | unknown |
| 328 | -2 | unknown |
| 390 | -2 | trypsin-like serine protease |
| 463 | -2 | ubiquitin-dependent endocytosis; arrestin |
| 484 | -2 | 7tm odorant receptor |

BLAST searches for the members of this gene family that were detected in *Ostrinia nubilalis* and *Manduca sexta* revealed very close (95-100%) matches to putative pheromone receptors in their respective species, indicating that this specific family of 7tm odorant receptors may be involved in pheromone recognition. Vogt *et al. (2015)* previously linked differences in mate recognition between butterflies and all other moths with the loss of a specific set of odorant binding proteins (OBPs), correlating said loss with a transition from scent-based to sight-based long distance mate recognition. Our finding of a significant pheromone receptor loss complements their hypothesis nicely, since OBPs facilitate the transport of molecules to odorant receptors. Similar trends have also been observed in primates, where the loss of function (pseudogenization) of odorant receptors has been linked with the acquisition of color vision (Gilad et al. 2004). Together, these results suggest a fundamental trade-off between investing in smell versus vision, and that patterns of odorant receptor loss may be common across Animalia upon shifts in the importance of visual information.

### Diet breadth is associated with the size of several important gene families

Our final analysis investigated whether the sizes of any gene families can be associated with species variation on a generalist/specialist axis (counts of the number of host orders used by each species are reported in Figure 1). Using the root-filtered, *de novo* protein set as input, we found that, of the 569 gene families included in our analysis, 12 were found to interact significantly with host order count (Fig. 4A; Table S9). Eight gene families were significantly larger with increasing diet breadth (Fig. 4B), including a family of carboxylesterases and a family of trypsin-like serine proteases. In contrast, four gene families were significantly larger with increasing diet specialization, including two chorion protein families. (Fig. 4C).

**Figure 4.**
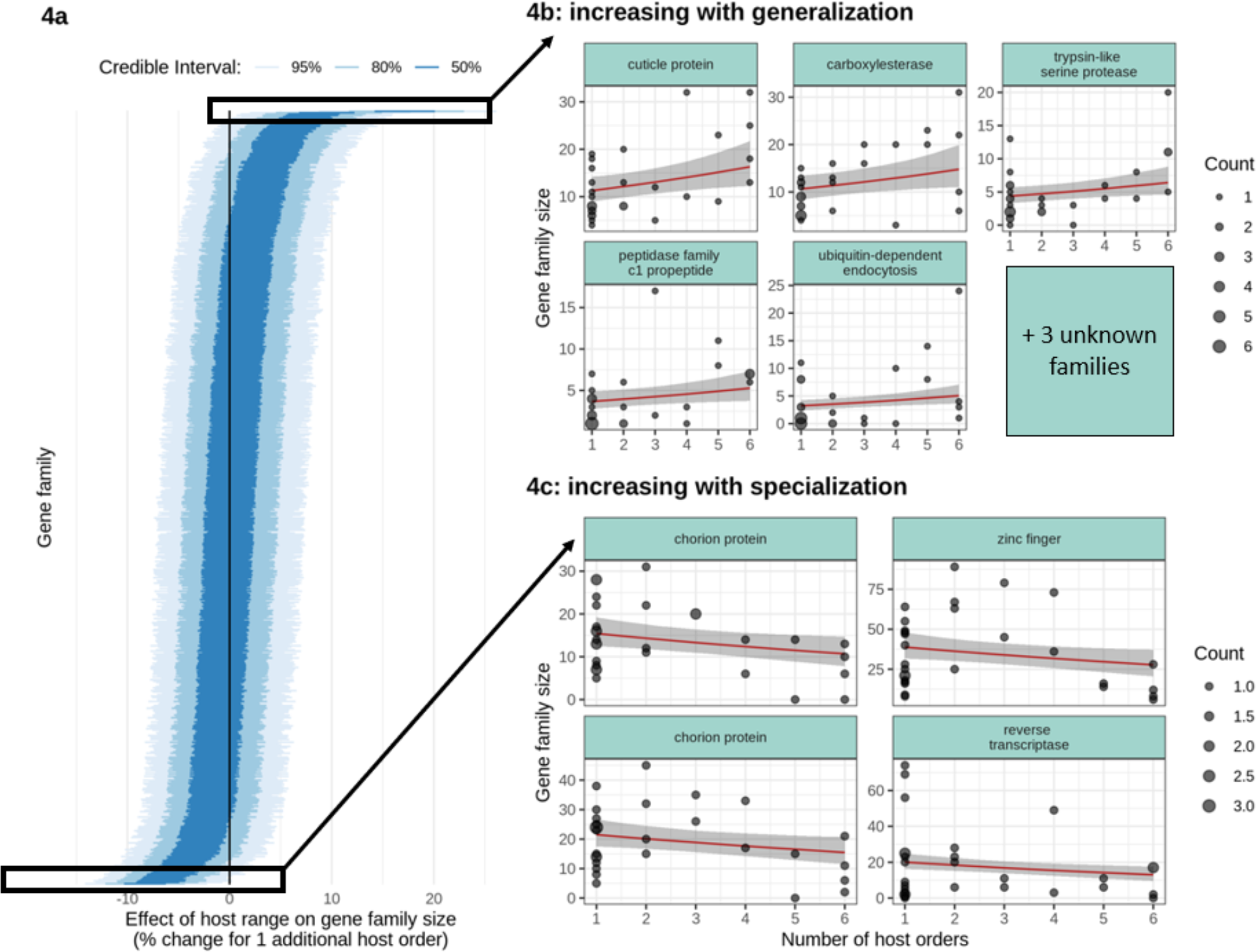
Effects of host range on gene family size. Host range size for each of n=30 Lepidoptera species was assessed by the number of orders larvae have been observed feeding on (see Table S7). (4A) The effect of host range is depicted as credible intervals (50%, 80% and 95%) of posterior distributions of slopes from a multilevel model (see Methods). Each horizontal line represents a gene family, and families are ordered by their posterior averages. Results are for all gene families analyzed in the root-filtered, lambda ≠ mu, BRAKER2 annotation set that had an average gene family size per species >= 3. The vast majority of the 569 gene families analyzed did not have their size significantly affected by host range, as seen by their overlap with 0 (solid line). Gene families that were significantly larger in species with more host orders (generalists) or significantly larger in species with fewer host orders (specialists) are shown in the insets 4B and 4C, respectively. Regression lines represent fits from the multilevel model, with the shaded region representing the 95% credible interval. As many species perfectly overlap, we scaled the point size to the number of species (‘Count’) at that coordinate.

Of the gene families that increased in size with increasing diet breath, the carboxylesterases and the serine proteases are perhaps the most relevant for feeding behavior, with the former being linked with general detoxification responses in insects (Devonshire 1977; Claudianos et al. 2006; Cao et al. 2008) and the latter found responsible for, e.g., immune responses and protein digestion (Zhu-Salzman & Zeng 2014; Cao et al. 2015; Jiang et al. 2010). For species encountering different classes of plant toxins, having several copies of these detoxification and digestion-related genes could prove advantageous (either due to increased protein production, or specialization of individual gene copies). Indeed, gene family expansions of serine proteases have been associated with the wide hostplant repertoire of the moth species *S. frugiperda*, with *S. frugiperda* being found to have 86 digestive serine protease genes compared to the 68 found in the specialist *M. sexta* (Gouin et al. 2017).

Our results for gene families that increased in size with increasing dietary specialization were harder to interpret, given the extreme variety found in chorion (i.e. egg casing) proteins (Kafatos et al. 1977). We speculate that an increase in chorion genes copy number could be an adaptation against plant defensive responses to Lepidoptera eggs, such as signaling for egg parasites (see, e.g., Tamiru et al. 2011; Cusumano et al. 2015; Hilker & Fatouros 2015) or hypersensitive response-like necrosis (described in Shapiro & DeVay 1987; Petzold-Maxwell et al. 2011; Fatouros et al. 2014; Griese et al. 2021). Although many of the mechanisms involved in plant-egg interactions are still poorly understood, it has been shown that hypersensitive response-like necrosis in cabbages is more severe when induced by eggs of butterflies specialized to cabbages, compared to eggs of non-specialized species (Griese et al. 2021). If specialist-targeted defenses are relatively common across the plant kingdom, it is possible that additional egg casing proteins are somehow beneficial to specialists.

Our analysis here is an improvement on previous associations of diet breadth and gene family sizes in Lepidoptera, as we used standardized, complete protein sets and accounted for the effects of phylogenetic relationships. Because we did not limit our study to consider only digestive and detoxification genes, we were able to identify surprising correlations between, e.g., diet breadth and copy numbers of egg casing genes. Furthermore, by factoring phylogenetic information into our model, we minimized the risk of phylogenetic pseudoreplication causing Type I error in our identification of significant correlations between diet breadth and the size of gene families (Middleton-Welling et al. 2020; Felsenstein 1985). This, along with differences in how genes were clustered, likely contributed to our not discovering significant correlations between increasing diet generalization and copy numbers of carboxyl- and choline esterase or glutathione-S-transferase genes, as was reported by Breeschoten et al. (2022).

### Where do we go now?

As demonstrated throughout this study, analyses of gene duplication and death are extremely useful tools for identifying candidate gene families of evolutionary interest. Our work advances this field by using a uniform set of annotated proteins for all genomes, investigating associations while correcting for phylogeny, and assessing all gene families rather than *a priori* subsets. Modifications of our model for correlating gene family sizes with diet breadth while taking into account phylogenetic relatedness can be used in future studies to correlate gene family size with other quantitative variables, such as habitat type or other aspects of life history. For butterfly and moth species, databases are growing for this type of trait information, such as the European and Maghreb Butterfly Trait Database (Middleton-Welling et al. 2020) and LepTraits (Shirey et al. 2022).

Fine-scale analyses (on the level of families or orders) will also soon be possible for many insect clades, as hundreds of new genomes will be released through projects like Darwin Tree of Life (https://www.darwintreeoflife.org/) in the coming years. Such analyses will be able to include far more gene families than we could in the present study, as the ability to detect gene families deeper through time will be increased by composed primarily of high quality genomes. Paired with information from trait databases and macroevolutionary studies it may be possible to use these fine-scale analyses to search for patterns of gene duplication or loss corresponding with repeated, yet evolutionarily independent, life history changes. When studying plant-insect interactions, one could explore if any genes tend to be duplicated or lost upon a host shift that has happened several times across a phylogeny and thereby identify candidate genes worthy of further study. For example, once candidate genes or gene families are identified by GBDD analyses, such as those used here, gene manipulations via RNAi or CRISPR-Cas9 could be used to assess their phenotypic effects, if such effects are not already documented. This will allow for functional testing of generated hypotheses, as well as for intersection with other datasets, such as those containing information on gene expression variation. In sum, although GBDD studies are demonstrably useful tools for generating evolutionary hypotheses, they are only a first step towards understanding how and why traits of interest emerged.

## Methods

### Genome datasets and assessment

Publicly available reference genomes for Lepidopteran species (n=30) were downloaded from LepBase (Challi et al. 2016) or NCBI (Table S1). Genome completeness was assessed using BUSCO scores (v. 4.1.2; Simão et al. 2015), with the OrthoDB v10 (Kriventseva et al. 2019) database of 5286 single-copy orthologs found among 16 Lepidoptera species, which allowed us to assess the presence, completeness and single copy status of these genes in each Lepidopteran genome we used. When available, genome annotations and protein sets provided with genomes were downloaded for use as native comparisons in downstream analyses (Table S1). For species that did not have genome annotations provided (see Table S2), or when we wanted to generate a uniform set of genome annotations for all our genomes, we generated genome annotations using the BRAKER2 pipeline, described below.

### Annotation generation & assessment

Because heterogeneity in annotation methods can inflate counts of lineage-specific genes in comparative analyses (Weisman et al. 2022), we began our main analyses of gene birth and death by generating annotations for all collected genomes with a standardized BRAKER2 pipeline. The BRAKER2 pipeline is an automated genome annotation pipeline that trains the gene prediction tools GeneMark-EX and Augustus, using either RNA-seq data from a target species or protein sets from closely related species (or both), and then uses the prediction tools to make gene predictions for an input genome (Stanke et al. 2006, 2008; Hoff et al. 2016, 2019; Brůna et al. 2021). Additionally, redundant training gene structures are filtered out with Diamond(Buchfink et al. 2015). In this study, we used the Arthropoda reference protein set from OrthoDB as our training dataset for gene model prediction (OrthoDB v10; Kriventseva et al. 2019), following the recommendation of BRAKER2. The steps in our BRAKER2 annotation pipeline were as follows. Genomes were first softmasked for repeat regions using redmask.py wrapper (v 0.0.2), which softmasks each genome with RED (v. 05/22/2015), a fully automated, self-training tool that can detect and mask both transposons and tandem repeats within the genome (Girgis 2015). After they were softmasked, genomes were used as input for BRAKER2 (using genome & protein mode), which generated gene annotations for each species. All protein sets per species used in this study (i.e. those that were downloaded, and those that were generated *de novo*) were filtered with the AGAT script keep_longest_isoform (Dainat 2021), so that only the longest isoform for each gene would be included in downstream analyses. Additionally, predicted proteins lacking start or stop codons, or with internal stop codons, were removed from the dataset. Following filtering, annotation quality was assessed with BUSCO scores (Simão et al. 2015; Manni et al. 2021) generated against the OrthoDB v10 Lepidoptera database (Kriventseva et al. 2019; Table S2).

### Comparison of native and BRAKER2 protein sets

To compare the content of native versus *de novo* annotation sets, we used the tool OrthoFinder (v. 2.5.1, Emms & Kelly 2019)) separately on the complete protein sets of each annotation type. This tool quickly assigned genes to orthogroups and then generated basic statistics on, e.g., the average number of genes per orthogroup. Output from the “Statistics_Overall” files from both OrthoFinder runs can be found in Table S6.

### Generating CAFE input

Two inputs were needed for our analysis of GBDD in CAFE (v 4.2.1; Han et al. 2013), which is software that explores gene family birth-death dynamics along a user-provided phylogenetic tree. These inputs were a clustering of all the protein datasets and a time-calibrated phylogenetic tree of all species. Following the CAFE recommendations, we clustered proteins into gene families across species with an all-against-all BLASTp (v. 2.5.0), using default settings. The resulting blast output was then clustered into “families” with the software mcl (v. 14-137; (van Dongen 2000; Enright et al. 2002) using the inflation value of 3 recommended in the CAFE tutorial. This pipeline of BLAST result clustering with mcl was performed separately for the native annotation set and the *de novo* annotation set.

The time tree for the species of Lepidoptera that have available reference genomes was reconstructed based on published studies. The relationships of the species and the times of divergence for most nodes were taken from Kawahara et al. (2019). Relationships and times of divergence within the genus *Papilio* were based on Allio et al. (2020), and within the genus *Heliconius* on Kozak et al. (2015).

### Estimating gene family dynamics

The clustered protein families (mcl output) were processed with CAFE, which, in addition to estimating counts of gene family members for each node, identifies gene families that are evolving rapidly (De Bie et al. 2006). For all CAFE runs in this study, we specified that gene families with an assigned p-value of 0.05 or less would be flagged as rapidly evolving. We chose settings where gene families that were assumed to not exist at the base of the phylogeny would be excluded (-filter flag), as not doing so artificially inflates CAFE’s gene death rate estimates. We excluded 27 mcl-defined gene families from our rate analyses due to their large size (we define a large family as one where one species has more than 100 gene copies), since large families are likely to cause non-informative estimations of lambda and mu if not removed (CAFE tutorial, 20 January 2016). These excluded families were later added in downstream analyses of diet breadth versus gene family size but were subsequently excluded again as they were dominated by transposable elements (Table S10).

In parallel with analyses of gene family expansions, CAFE was also used to calculate gene duplication rate (ƛ) as a global value across the tree, where birth rate was assumed equal to death rate (μ). We also analyzed a more complex model where gene death rate (μ) was not assumed to be equal to ƛ. Models of the same annotation type were assessed for fit with their model score (a negative log likelihood value), where scores closer to zero indicated a better fit.

We accounted for error in the assemblies and annotations of the input data by running CAFE several times with the same input data, but with a different error distribution for each run (cafeerror.py script). The error distribution with the lowest log likelihood score was then used as an input (-errormodel flag) in future CAFE runs to give less biased estimations of both gene counts and gene duplication-death rates at each node.

To visualize differences in rates of gene births and deaths across the phylogeny, information from CAFE output files was overlaid on our species tree with the tidyverse (v 1.3.0; Wickham et al. 2019), phytools (v 0.7.70; Revell 2012)), and ggtree (v. 2.2.4; Yu et al. 2017, 2018; Yu 2020) packages for the statistical software R (v. 4.0.1, R Core Team 2020). Each of the R scripts used for plotting the results of this study can be found at https://github.com/pierid-yay/Lep_Diet_Breadth_Evolution_2021. We also provide a flowchart of our process for generating CAFE input files in Figure S2.

### Gene family functional annotation analyses

For all gene families that were deemed rapidly evolving (p < 0.05) by CAFE at the crown of Papilionoidea, gene sequences were extracted from the appropriate protein datasets and functionally annotated with the software eggNOG-mapper (v. 5.0; Huerta-Cepas et al. 2017, 2019)

### Assessing gene family size in relation to hostplant feeding range

To test whether the size of gene families and host breadth are correlated along the Lepidoptera phylogeny, we modeled the per family relationship between gene count and number of host plant orders in a single multilevel Bayesian model. We estimated the number of hostplant orders used by each species by reviewing hostplant records on FUNET (Savela 2021) and checking them against other web and literature sources (see Figure 1 for counts and Table S8 for references). We included only commonly used hosts supported by several sources in order to avoid erroneous records. Our analysis then considered the size of each gene family per species as the response variable, modeled using the negative binomial distribution with a log link function. To reduce the computational effort of fitting this model we excluded very small gene families (mean gene count per species < 3), as we would have very limited power to detect significant effects in this group. As population-level (or fixed) effects we included an intercept and the effect of hostplant order count. Additionally, we added group-level (or random) coefficients for the intercept and the effect of hostplant order count per gene family. To account for the non-independence due to multiple observations of the same species and due to phylogenetic inertia, we included two group-level effects for species, uncorrelated varying intercepts and varying intercepts correlated across the phylogenetic covariance matrix.

For inference, we obtained posterior distributions of the slope between the number of genes and the number of hostplant orders from this model, for each gene family. We expressed these slopes as incidence ratios for ease of interpretation. We classified those gene families where the 95% credible intervals did not overlap zero as having a clearly identifiable and significant relationship with host breadth. Note that these slope estimates have undergone shrinkage towards the mean slope across gene families, and we therefore do not perform further adjustments for multiple testing.

Parameter values were estimated using the brms (Bürkner 2017, 2018) interface to the probabilistic programming language Stan (Carpenter et al. 2017).We used weakly informative prior distributions, with a StudentT(3, 0, 1) prior for the population level slope. For the group level standard deviations and residual standard deviation, we used the positive range of StudentT(3, 0, 2.5). The shape parameter used Gamma (0.01, 0.01). Finally, we used an LKJ (1) prior for the group-level correlation between intercept and slope, which is uniform over the range −1 to 1. Posterior distributions were obtained using Stan’s no-U-turn HMC sampler, with 4 chains of 3000 iterations, with the first 1000 used as warm-up and discarded. We additionally set the adapt delta parameter to 0.9. We evaluated model fit using posterior predictive plots.

For all instances where we identified a significant effect of hostplant order count on gene family size, we generated functional annotations for the relevant gene family with eggNOG-mapper v. 5.0 (Huerta-Cepas et al. 2019)

## Data Accessibility Statement

Script examples and files needed to replicate this study are available at the first author’s github page (https://github.com/pierid-yay/Lep_Diet_Breadth_Evolution_2021) or in previously published studies. A full set of output files will be uploaded to Dryad upon acceptance.

