## Supplemental for "Genome-wide gene birth-death dynamics are associated with diet breadth variation in Lepidoptera"

### Contents


Supplementary Fig. 1: Counts of CAFE-estimated gene family expansions and contractions across the Lepidoptera phylogeny

Supplementary Fig 2: Flowchart of CAFE analysis workflow

Table legends for supplementary tables
(found in file Dort_2022_SuppTables.xlsx)

Supplementary references

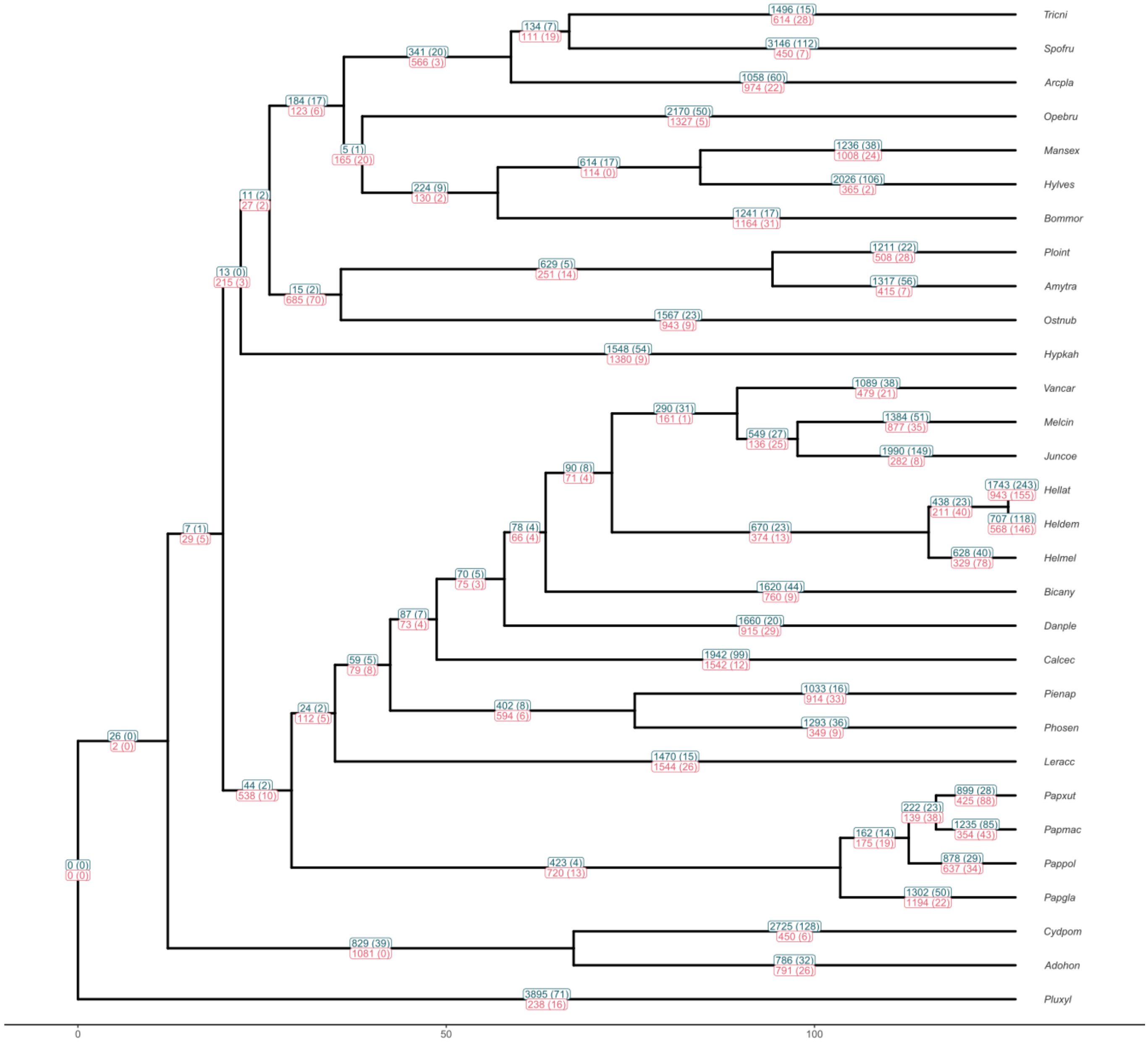


**Supplementary Figure 1:** Counts of CAFE-estimated gene family expansions (in green) and contractions (in red) for filtered BRAKER2 protein sets. Numbers in parentheses indicate the count of significant ($p < 0.05$) expansions or contractions at each node.

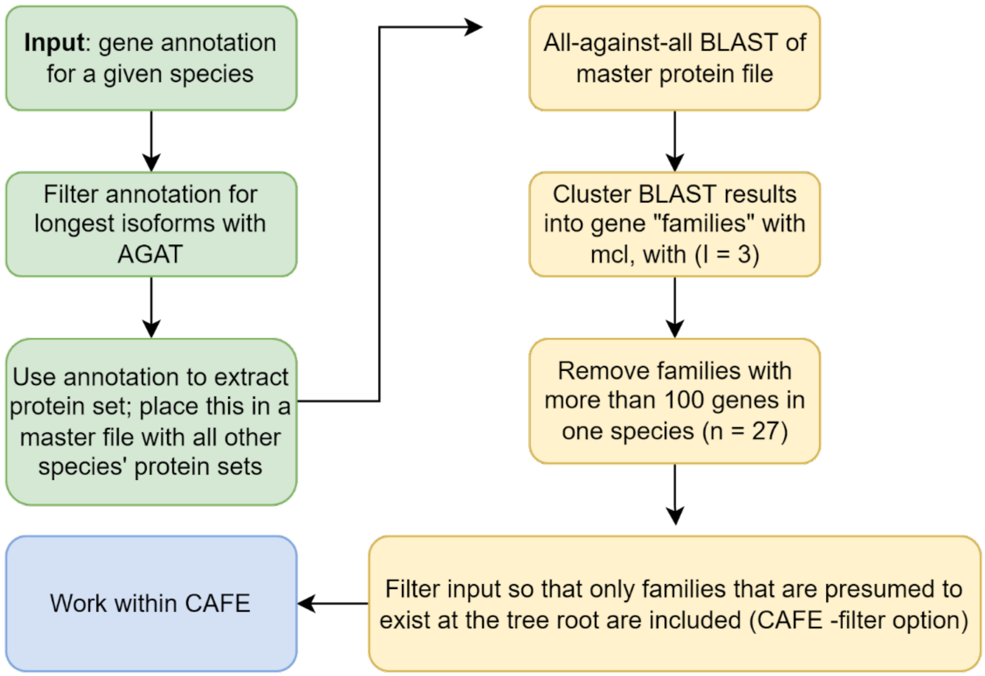


**Supplementary Figure 2:** Generation of input files for gene birth-death analysis of protein datasets. The above workflow was used for this study's *de novo* annotation set of 30 Lepidopteran genomes.

**Table legends for supplementary tables (found as tabs in Dort_2022_SuppTables.xlsx)**

**Supplementary Table 1**: Species information, reference genome and native annotation metrics, and BUSCO scores. Taxonomic information for all species (n = 30) included in this study is presented here, along with information on where their respective reference genomes were obtained. Basic metrics for reference genomes and native annotations (e.g. genome size, genome N50, protein counts) are also reported, with "native_protein_fasta_count" values referring to the number of proteins in the raw (unfiltered) protein fasta files provided with assembly downloads . BUSCO scores for downloaded genomes are reported in the furthest right columns as a measure of genome completeness.

**Supplementary Table 2:** Gene counts for all annotations generated with the BRAKER2 pipeline described in the Methods section of this study. Counts after isoform filtering with AGAT and further filtering to only include genes with correct start and stop codons are reported in the columns "Isoform_filtered_b2_count" and "Isoform_startstop_clean_b2_count", respectively. In some cases, "native" annotations were generated with BRAKER2 and were therefore identical to braker2 annotations. These genomes are marked with a "Y" in the column labelled "Orig_w_same_pipeline".

**Supplementary Table 3**: Per-species gene counts and BUSCO scores for filtered, BRAKER2-generated *de novo* protein sets.

**Supplementary Table 4:** BUSCO scores for the native set of protein fastas provided with genome downloads.

**Supplementary Table 5:** Gene counts generated from native annotation (.gff) files both before and after filtering steps. Percent decrease in gene count between raw and "start-stop" filtered annotations is also reported. Native annotation files were available for 19 of 30 species in this study; for species where a native annotation did not exist, a value of "DNE" was entered in the native_gff column.

**Supplementary Table 6:** Expanded OrthoFinder results for both native and *de novo* protein sets. All values reported here originate from the OrthoFinder-generated "Statistics_Overall.tsv" files.

**Supplementary Table 7:** Full set of model scores and birth/death rates from CAFE GBDD analyses, using filtered *de novo* protein sets as input. Average birth and death rates across runs are also calculated and reported here.

**Supplementary Table 8:** Hostplant order information. Here, we list the hostplant order counts and identities for each species in this study. For information not obtained from FUNET, relevant citations are listed under "Source".

**Supplementary Table 9:** Gene families identified as either significantly increasing or decreasing in size with increases in diet breadth. A value of "S" in the "Increases_with" column denotes that the family increased in size with increasing diet specialization. A value of "G" denotes that the family increased in size with increasing diet generalization. Functional annotations in the “Annotation” column were generated with eggNOG, as reported in the Methods section of this study.

**Supplementary Table 10:** Results from a run of our diet breadth vs. per-family gene count analyses where we included the 27 large gene families excluded from gene duplication and death rate estimates (see Methods).

Kanost, M. R., E. L. Arrese, X. Cao, Y.-R. Chen, S. Chellapilla, M. R. Goldsmith, E. Grosse-Wilde, D. G. Heckel, N. Herndon, H. Jiang, A. Papanicolaou, J. Qu, J. L. Soulages, H. Vogel, J. Walters, R. M. Waterhouse, S.-J. Ahn, F. C. Almeida, C. An, P. Aqrawi, A. Bretschneider, W. B. Bryant, S. Bucks, H. Chao, G. Chevignon, J. M. Christen, D. F. Clarke, N. T. Dittmer, L. C. F. Ferguson, S. Garavelou, K. H. J. Gordon, R. T. Gunaratna, Y. Han, F. Hauser, Y. He, H. Heidel-Fischer, A. Hirsh, Y. Hu, H. Jiang, D. Kalra, C. Klinner, C. König, C. Kovar, A. R. Kroll, S. S. Kuwar, S. L. Lee, R. Lehman, K. Li, Z. Li, H. Liang, S. Lovelace, Z. Lu, J. H. Mansfield, K. J. McCulloch, T. Mathew, B. Morton, D. M. Muzny, D. Neunemann, F. Ongeri, Y. Pauchet, L.-L. Pu, I. Pyrousis, X.-J. Rao, A. Redding, C. Roesel, A. Sanchez-Gracia, S. Schaack, A. Shukla, G. Tetreau, Y. Wang, G.-H. Xiong, W. Traut, T. K. Walsh, K. C. Worley, D. Wu, W. Wu, Y.-Q. Wu, X. Zhang, Z. Zou, H. Zucker, A. D. Briscoe, T. Burmester, R. J. Clem, R. Feyereisen, C. J. P. Grimmelikhuijzen, S. J. Hamodrakas, B. S. Hansson, E. Huguet, L. S. Jermiin, Q. Lan, H. K. Lehman, M. Lorenzen, H. Merzendorfer, I. Michalopoulos, D. B. Morton, S. Muthukrishnan, J. G. Oakeshott, W. Palmer, Y. Park, A. L. Passarelli, J. Rozas, L. M. Schwartz, W. Smith, A. Southgate, A. Vilcinskas, R. Vogt, P. Wang, J. Werren, X.-Q. Yu, J.-J. Zhou, S. J. Brown, S. E. Scherer, S. Richards, and G. W. Blissard. 2016. Multifaceted biological insights from a draft genome sequence of the tobacco hornworm moth, Manduca sexta. Insect Biochem Mol Biol 76:118–147.

Lara-Villalón, M., V. Vanoye-Eligio, M. A. Solís, G. Sánchez-Ramos, and J. C. Chacón-Hernández. 2017. The Navel Orangeworm, Amyelois transitella (Walker) (Lepidoptera: Pyralidae), Discovered in Northeastern Mexico Feeding on Sapindaceae. went 119:601–605. Entomological Society of Washington.

Lewis, J. J., K. R. L. van der Burg, A. Mazo-Vargas, and R. D. Reed. 2016. ChIP-Seq-Annotated Heliconius erato Genome Highlights Patterns of cis-Regulatory Evolution in Lepidoptera. Cell Rep 16:2855–2863.

Li, X., D. Fan, W. Zhang, G. Liu, L. Zhang, L. Zhao, X. Fang, L. Chen, Y. Dong, Y. Chen, Y. Ding, R. Zhao, M. Feng, Y. Zhu, Y. Feng, X. Jiang, D. Zhu, H. Xiang, X. Feng, S. Li, J. Wang, G. Zhang, M. R. Kronforst, and W. Wang. 2015. Outbred genome sequencing and CRISPR/Cas9 gene editing in butterflies. Nat Commun 6:8212. Nature Publishing Group.

Mercader, R. J., M. L. Aardema, and J. M. Scriber. 2009. Hybridization Leads to Host-Use Divergence in a Polyphagous Butterfly Sibling Species Pair. Oecologia 158:651–662. Springer, International Association for Ecology.

Nadeau, N. J., M. Ruiz, P. Salazar, B. Counterman, J. A. Medina, H. Ortiz-Zuazaga, A. Morrison, W. O. McMillan, C. D. Jiggins, and R. Papa. 2014. Population genomics of parallel hybrid zones in the mimetic butterflies, H. melpomene and H. erato. Genome Res 24:1316–1333.

Nishikawa, H., T. Iijima, R. Kajitani, J. Yamaguchi, T. Ando, Y. Suzuki, S. Sugano, A. Fujiyama, S. Kosugi, H. Hirakawa, S. Tabata, K. Ozaki, H. Morimoto, K. Ihara, M. Obara, H. Hori, T. Itoh, and H. Fujiwara. 2015. A genetic mechanism for female-limited Batesian mimicry in Papilio butterfly. Nat Genet 47:405–409. Nature Publishing Group.

Nowell, R. W., B. Elsworth, V. Oostra, B. J. Zwaan, C. W. Wheat, M. Saastamoinen, I. J. Saccheri, A. E. van't Hof, B. R. Wasik, H. Connahs, M. L. Aslam, S. Kumar, R. J. Challis, A. Monteiro, P. M. Brakefield, and M. Blaxter. 2017. A high-coverage draft genome of the mycalesine butterfly Bicyclus anynana. GigaScience 6.

Ojala, K., R. Julkunen-Tiitto, L. Lindström, and J. Mappes. 2005. Diet affects the immune defence and life-history traits of an Arctiid moth Parasemia plantaginis. Evolutionary Ecology Research 7:1153–1170.

Pippel, M., D. Jebb, F. Patzold, S. Winkler, H. Vogel, G. Myers, M. Hiller, and A. K. Hundsdoerfer. 2020. A highly contiguous genome assembly of the bat hawkmoth Hyles vespertilio (Lepidoptera: Sphingidae). Gigascience 9.

Schmitz, P., and D. Rubinoff. 2011. The Hawaiian amphibious caterpillar guild: new species of Hyposmocoma (Lepidoptera: Cosmopterigidae) confirm distinct aquatic invasions and complex speciation patterns. Zoological Journal of the Linnean Society 162:15–42.

Shi, B.-C., W. Liu, and S.-J. Wei. 2013. The complete mitochondrial genome of the codling moth Cydia pomonella (Lepidoptera: Tortricidae). Mitochondrial DNA 24:37–39.

Uchibori-Asano, M., A. Jouraku, T. Uchiyama, K. Yokoi, G. Akiduki, Y. Suetsugu, T. Kobayashi, A. Ozawa, S. Minami, C. Ishizuka, Y. Nakagawa, T. Daimon, and T. Shinoda. 2019. Genome-wide Identification of Tebufenozide Resistant Genes in the smaller tea tortrix, Adoxophyes honmai (Lepidoptera: Tortricidae). Sci Rep 9:4203.

van der Burg, K. R. L., J. J. Lewis, A. Martin, H. F. Nijhout, C. G. Danko, and R. D. Reed. 2019. Contrasting Roles of Transcription Factors Spineless and EcR in the Highly Dynamic Chromatin Landscape of Butterfly Wing Metamorphosis. Cell Rep 27:1027-1038.e3.

Yen, E. C., S. A. McCarthy, J. A. Galarza, T. N. Generalovic, S. Pelan, P. Nguyen, J. I. Meier, I. A. Warren, J. Mappes, R. Durbin, and C. D. Jiggins. 2020. A haplotype-resolved, de novo genome assembly for the wood tiger moth (Arctia plantaginis) through trio binning. Gigascience 9.

You, M., Z. Yue, W. He, X. Yang, G. Yang, M. Xie, D. Zhan, S. W. Baxter, L. Vasseur, G. M. Gurr, C. J. Douglas, J. Bai, P. Wang, K. Cui, S. Huang, X. Li, Q. Zhou, Z. Wu, Q. Chen, C. Liu, B. Wang, X. Li, X. Xu, C. Lu, M. Hu, J. W. Davey, S. M. Smith, M. Chen, X. Xia, W. Tang, F. Ke, D. Zheng, Y. Hu, F. Song, Y. You, X. Ma, L. Peng, Y. Zheng, Y. Liang, Y. Chen, L. Yu, Y. Zhang, Y. Liu, G. Li, L. Fang, J. Li, X. Zhou, Y. Luo, C. Gou, J. Wang, J. Wang, H. Yang, and J.

Wang. 2013. A heterozygous moth genome provides insights into herbivory and detoxification. Nat Genet 45:220–225. Nature Publishing Group.

Zhan, S., W. Zhang, K. Niitepõld, J. Hsu, J. F. Haeger, M. P. Zalucki, S. Altizer, J. C. de Roode, S. M. Reppert, and M. R. Kronforst. 2014. The genetics of monarch butterfly migration and warning coloration. Nature 514:317–321.
